# Constructing a holistic map of cell fate decision by hyper solution landscape

**DOI:** 10.1101/2024.11.28.625944

**Authors:** Xiaoyi Zhang, Zhiyuan Li, Lei Zhang

## Abstract

The Waddington landscape metaphor has inspired extensive quantitative studies of cell fate decisions using dynamical systems. While these approaches provide valuable insights, the intrinsic nonlinear complexity and the parameter dependence limits systematic analysis of fate transitions. Here, we introduce the Hyper Solution Landscape (HSL), a minimally parameter-dependent methodology showing a comprehensive structure of all possible landscape configurations for gene regulatory networks. HSL connects different solution landscapes to reflect dynamic changes of the landscapes associated with bifurcations. Applied to the Cross-Inhibition with Self-activation motif, HSL analysis identifies key hyperparameters driving distinct directional changes in cell fate propensity. Different routes through the HSL between the same initial and final states can produce markedly different fate distributions. This enables rational design of transition strategies. We validate HSL’s utility in the seesaw model of cellular reprogramming, establishing a powerful framework for understanding and engineering cell fate decisions. A record of this paper’s Transparent Peer Review process is included in the Supplemental Information.

## Introduction

The Waddington landscape has served as a powerful metaphor for understanding cell fate decisions since its inception^1^, catalyzing extensive research in developmental biology^2,3^. Following Kauffman’s pioneering work using Boolean networks to characterize the valleys of this landscape^4,5^, researchers have made significant strides in quantifying cell fate decisions through dynamical systems, particularly gene regulatory networks (GRNs) ^6–9^. This quantitative framework has established a crucial link between mathematical attractors and biological cell fates, enabling computational studies of cellular decision-making through nonlinear dynamics^10–12^.

Among the various regulatory motifs studied, the Cross-Inhibition with Self-activation (CIS) pattern has emerged as particularly significant^13–17^, appearing in numerous biological contexts including GATA1-PU.1^18^, OCT4-CDX2^15^, and NANOG-GATA6 interactions^19^. Cell state transitions within these networks can occur through two primary mechanisms: stochastic fluctuations (noise-driven)^20–23^ or changes in external signals (signal-driven)^24–28^. These biological processes correspond to static and dynamic landscapes, respectively^29^.

While nonlinear dynamics models have proven valuable in characterizing cell fate decisions, they face a fundamental limitation: parameter dependency. Recent advances in nonequilibrium statistical physics have enabled the construction of energy landscapes from dynamical systems^30–34^. However, experimental determination of all relevant parameters in a gene regulatory network remains challenging, if not impossible, and different parameter sets can lead to distinct landscapes with varying predictions on cell fate behaviors^35^. Furthermore, the complexity of energy landscapes, which contain numerous details beyond the primary attractors, makes systematic analysis of global changes particularly challenging. While recent geometric approaches to cell fate decisions have provided valuable insights^36–39^, their abstract, gene-free nature has limited their direct application to real genetic networks^40^.

The solution landscape concept offers a promising alternative approach^41^. This framework creates a pathway map comprising all stationary points and their connections^42^, having demonstrated utility across various fields including gene regulatory networks^43^, liquid crystals^44^, and quasicrystals^45^. Unlike traditional energy landscapes that expend computational resources on fine details, solution landscapes focus on the essential elements of noise-driven cell fate decisions: the attractors (stable cell fates), saddle points (unstable states, as approximations to potential transition states in non-equilibrium systems^46^, forming a structural backbone upon which stochastic trajectories) and their connections (transition pathways). Their simplified graph structure of nodes and edges provides an efficient framework for analyzing static landscape and transitions driven by stochastic fluctuations. However, experimental manipulation of cell fates typically falls into the signal-driven category, where external signals alter the landscape geometry^43,47^. In these cases, the solution landscape itself undergoes transformation as external conditions change^48^. This dynamic nature calls for a more systematic approach to connect and analyze different solution landscapes, enabling better understanding and control of directed cell fate decisions^49^.

Building on this foundation, we developed a framework: Hyper Solution Landscape (HSL), which is the ensemble of all solution landscapes and their bifurcation relationships for a dynamic system. HSL is a minimally parameter-dependent methodology that shows a comprehensive structure of all possible landscape configurations from a given GRN topology. By connecting different solution landscapes through bifurcation relationships, HSL provides a unified analysis for both noise-driven and signal-driven cell fate decisions. We apply this approach to both the CIS motif of differentiation and the seesaw model of reprogramming, establishing clear connections between solution landscape alterations and the Waddington landscape. Our analysis reveals intrinsic nonlinear complexity of cell fate decisions, including the most probable fate distributions and their responses to varying hyperparameters. Furthermore, we demonstrate how different bifurcation routes lead to distinct cell fate behaviors, enabling the rational design of cell fate decision routes through HSL.

## Results

### Solution Landscape Depicts the Geometry of Cell Fate Decisions Under a Given Signal Level

The dynamics of cell fate decisions are often studied through mathematical models of gene regulatory networks. Among these, the Cross-Inhibition with Self-activation (CIS) motif has emerged as a particularly important regulatory structure^13^ (Figure 1A). In this motif, two lineage-specifying transcription factors interact: factor X promotes its own expression while inhibiting factor Y, and vice versa. For such combinatory regulation^43^, the Shea-Ackers formalism, which mechanistically represents probabilistic occupancy of transcription factor binding sites on promoters^50,51^, provides broader coverage of regulatory logics than manual combinations of Hill functions(e.g., summation or multiplication)^30^, avoiding arbitrary choices that can influence outcomes. Therefore, we unified diverse regulatory scenarios into this flexible framework:

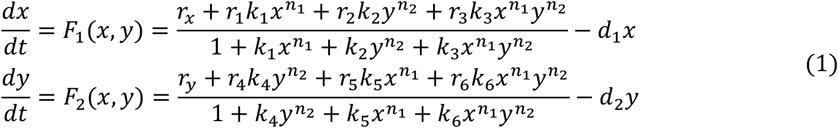

*n*_1_and *n*_2_ represent transcription factor cooperation efficiencies, *d*_1_ and *d*_2_ are degradation rates, *k*_1_ − *k*_6_ denote relative binding strengths to cis-regulatory elements, and *r*_*x*_, *r*_*y*_, *r*_1_ − *r*_6_, represent transcription rates for different binding configurations. For analytical convenience, we can express this system in vector form:

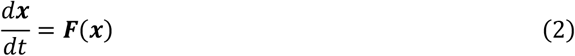

where the ***x*** = (*x, y*) and ***F***(***x***) = [*F*_1_(*x, y*), *F*_2_(*x, y*)].

**Figure 1.**
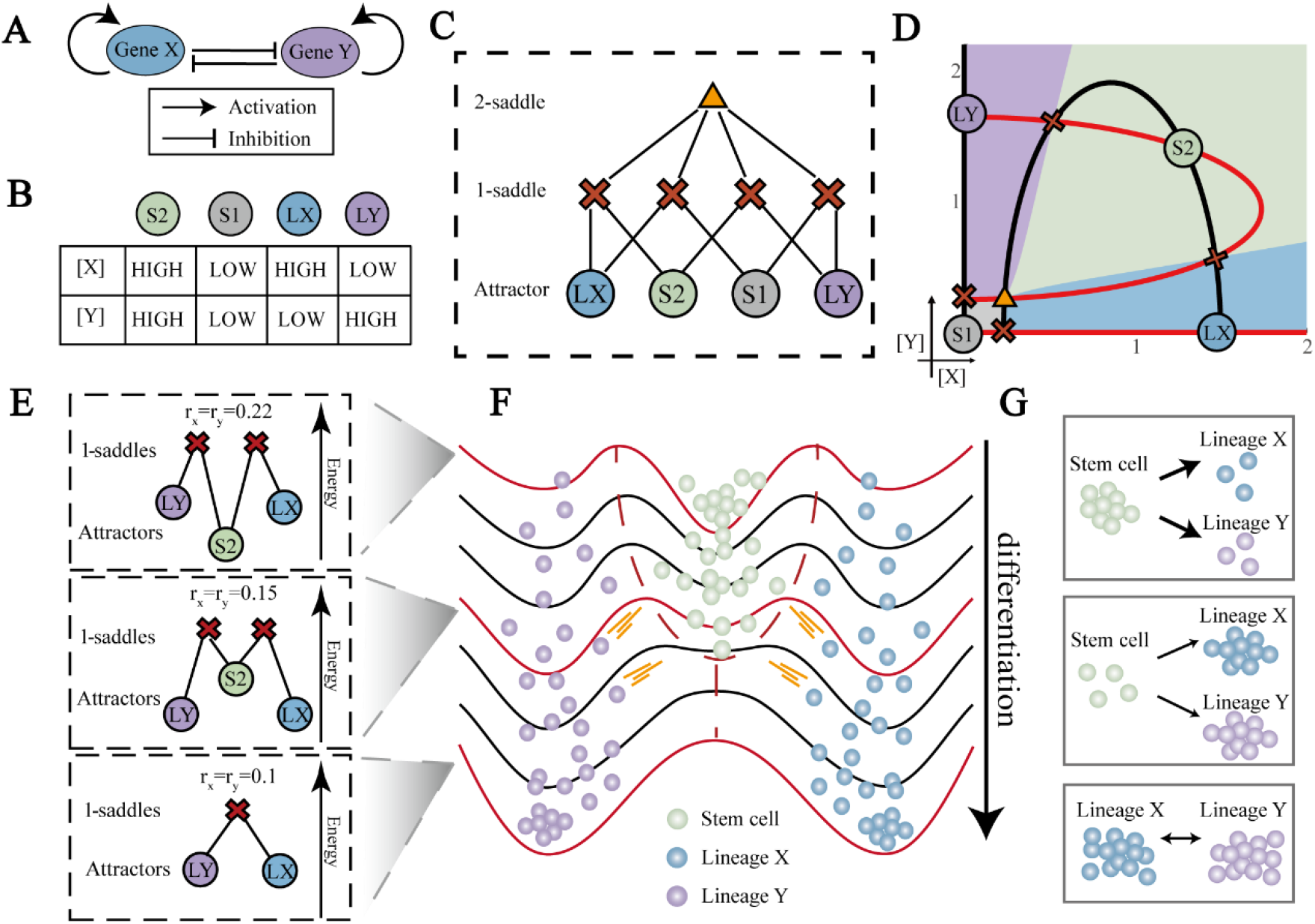
Solution landscape characterizes the geometry of cell fate decisions. (A) The cross-inhibition and self-activation (CIS) gene regulatory motif in cell fate decisions. Gene X and Y represent two antagonizing lineage-specifying genes. (B) Relationship between cell fates (i.e. attractors) and gene expression levels for the model in (A). Symbols S2, S1, LX, and LY represent uncommitted progenitor cell fates with high and low lineage specifying genes, lineage X, and lineage Y cell fates, respectively. These symbols are used for all following figures. (C) A solution landscape of the CIS motif in (A). The lowest layer of the solution landscape is composed of stable attractors represented by colored circles, which are labeled as LX, S2, S1, and LY to represent corresponding cell fates. The middle layer is composed of four index-1 saddle points represented by red crosses, and the top layer contains an index-2 saddle point represented by yellow triangle. The line in the figure indicates the connection between these points. These annotations are used for the following figures. (D) The phase diagram of the CIS motif in (A). The coordinate axis represents the expression of X and Y genes, respectively. The black line and the red line represent nullclines of 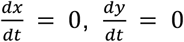, respectively. The intersections of nullclines are the stationary points in the system, with the same symbol as that in (C). Basins of attractors are filled by their corresponding colors. These annotations are used for the following figures. (E) Three solution landscapes during the signal-driven differentiation from uncommitted progenitor cell to Lineage X and Lineage Y, corresponding to the red lines in (F). The height of each point is determined by the quasi-potential. The parameters (*r*_*x*_, *r*_*y*_) corresponding to different solution landscapes are shown in the figure. (F) The schematic Waddington landscapes for the differentiation shown in (E). In each curve, the valleys represent attractors, and the blue, green and purple balls represent cells with LX, S2 and LY fates, respectively. The red dashed line indicates the boundary of the S2 attractor. The situation corresponding to the red line is shown in (E) and (G). (G) The schematic representation of cell fate decisions under three different situations, as corresponds in (E).

In the CIS motif, four distinct cell fates emerge as stable gene expression patterns, corresponding to mathematical attractors (Figure 1B). These states mirror the ‘valleys’ in Waddington’s landscape metaphor: two uncommitted progenitor cell states—S1 (low expression of both X and Y) and S2 (high expression of both)—and two lineage-committed states—LX (high X, low Y) and LY (low X, high Y). Between these stable states lie saddle points, analogous to the “peaks” in Waddington’s landscape, which separate different attractor basins. Of particular importance are the index-2 saddle points, which connect multiple index-1 saddle points and help define attractor basin boundaries.

By connecting all stationary points—both attractors and saddles—the solution landscape provides a complete geometric representation of all possible cell fate decisions under specific conditions^41^. The solution landscape is a hierarchical structure with all stationary points in different layers (Figure 1C). The General High-index Saddle Dynamics (GHiSD) method allows us to efficiently compute the index-k saddle points ^42^:

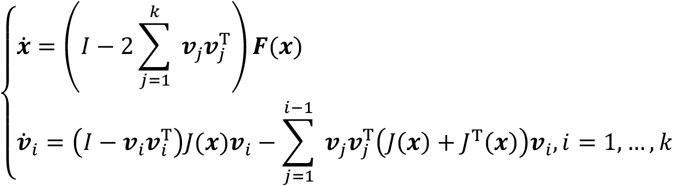

Where *J*(***x***) = ∇***F***(***x***) is the Jacobian matrix of ***F***(***x***) and the *v* is the eigenvector of *J*(***x***) corresponding to the positive eigenvalues.

The solution landscape reveals a static landscape under a given signal. For our two-dimensional (2D) system, the highest index for saddles is 2, and the solution landscape can be constructed hierarchically from the index-2 saddle point (repeller) down to the attractors using GHiSD combined with downward/upward searching algorithms^42^. The resulting landscape for the CIS motif reveals a single index-2 saddle point, four index-1 saddle points, and four stable attractors, arranged in a hierarchical structure that corresponds to the intersections of nullclines in state space (Figure 1D). By calculating the energy barrier (i.e. quasi-potential) between different attractors by the Geometric Minimum Action Method (GMAM)^52,53^ (Figure 1E and Methods), the solution landscape offers a perspective of noise-driven cell fate decision.

### The Alteration of Solution Landscapes Corresponds to Distinct Geometry of Cell Fate Decision

A solution landscape captures the complete set of stationary points—both saddles and attractors—under specific parameter conditions of a system, representing all possible cell fates and their transition relationships in a static landscape. In biological systems, varying signaling strengths manifest as parameter changes in mathematical models. These parameter shifts can alter the solution landscape, leading to changes in both the number of stable cell fates and the likelihood of transitions between them.

Consider the signal-induced differentiation process from uncommitted progenitor cells (S2) to lineage-specified cells (LX and LY) as an illustrative example. This process demonstrates how parameter-induced changes in the solution landscape govern the geometry of cell fate decisions: Initially, the solution landscape exhibits three stable attractors connected by two index-1 saddle points (Figure 1E, top panel). This configuration represents a uncommitted cell fate (S2) positioned between two differentiated states (LX and LY). As the parameter *r*_*x*_ and *r*_*y*_ decreases from 0.22 to 0.15 (Figure 1E, middle panel), the landscape’s topology remains unchanged, but the relative stabilities shift : the uncommitted progenitor cell state S2 becomes less stable, while the differentiated states LX and LY gain more stability. Further reduction of *r*_*x*_ and *r*_*y*_ to 0.1 triggers a subcritical fork bifurcation, causing the stable attractor corresponding to the uncommitted progenitor cell state to vanish, leaving only the LX and LY attractors connected by a saddle point (Figure 1E, bottom panel).

These progressive changes in the solution landscape directly correspond to cellular behavior. In the initial configuration, uncommitted progenitor cells predominantly maintain their undifferentiated state due to the higher stability of the S2 attractor (Figure 1G, top panel). As *r*_*x*_ and *r*_*y*_ decreases, the reduced stability of the uncommitted progenitor cell fate increases the probability of spontaneous differentiation (Figure 1G, middle panel). Following the bifurcation, the topological transformation of the landscape drives all uncommitted progenitor cells toward differentiated fates (Figure 1G, bottom panel).

This analysis reveals how signal-induced changes in cell fate decisions manifest as geometric transformations in solution landscapes. Bifurcations create qualitative changes in cell fate geometries, highlighting the need for tools capable of characterizing sequential changes in solution landscapes. Such tools would enable systematic analysis of complex cell fate transition processes.

### Hyper Solution Landscape (HSL): Ensemble of Solution Landscapes and Bifurcations

In order to gain a holistic view of cell fate decisions, we introduce the HSL, defined as the ensemble of solution landscapes and their bifurcation relationships, for a given gene regulatory network across all parameter sets. For the CIS motif, the HSL takes the form of a hierarchical graph (Figure 2A), with nodes representing topologies of solution landscapes and edges representing bifurcations. The nodes are arranged horizontally based on their number of stationary points and assigned unique identifiers based on their position in the graph. To facilitate readers’ tracking of the CIS motif, every node also has a descriptive name based on the number of saddle points and the types of attractors (Methods).

**Figure 2.**
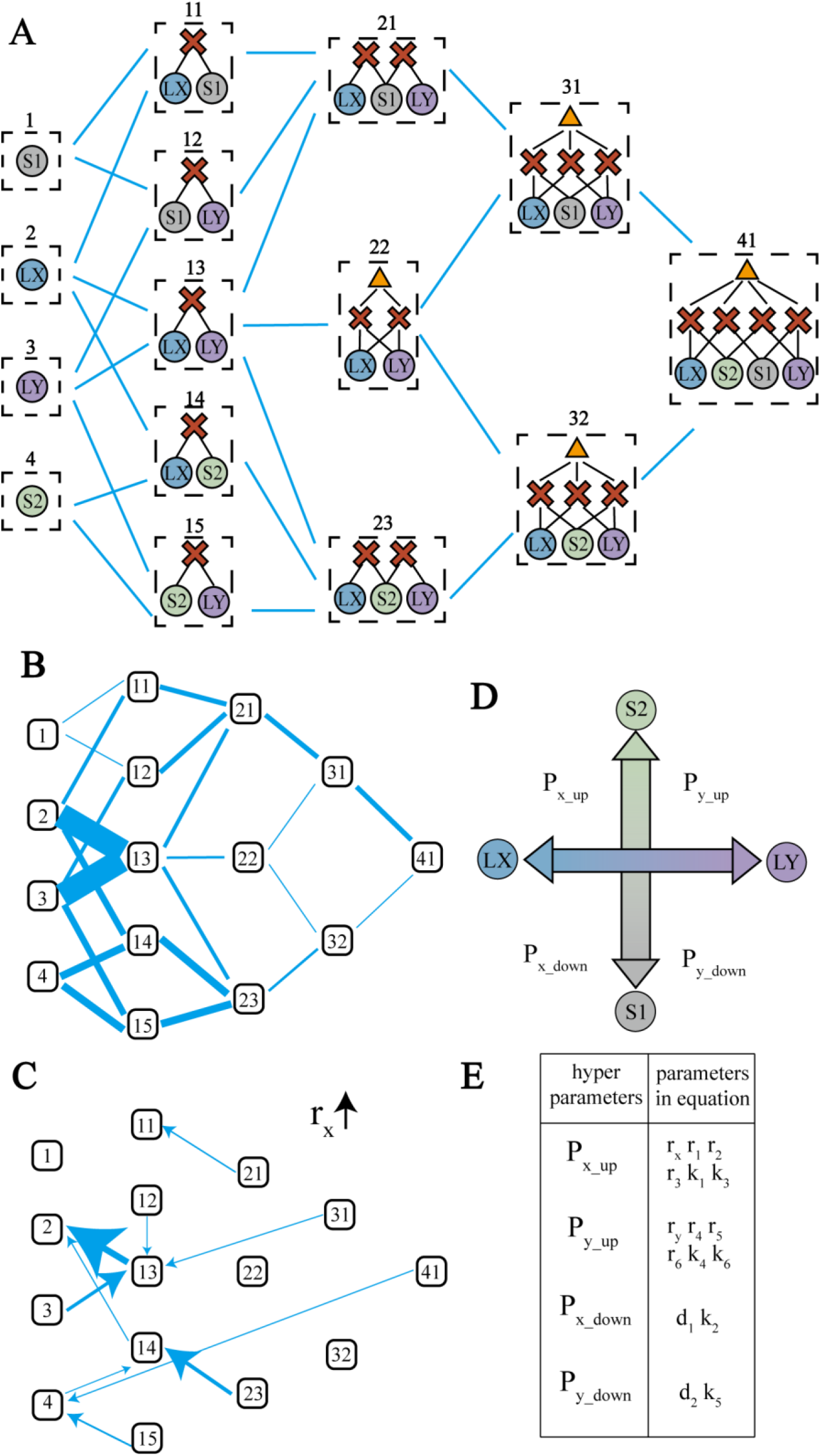
HSL connects different solution landscape topologies by bifurcations. (A) The HSL for the CIS motif shown in Figure 1(A). Individual solution landscape topology in each dotted box represents the node of HSL, and their IDs are shown above each box. Bifurcations induced by single parameter between solution landscape topologies are edges of HSL. (B) The HSL with weighted edges. Each solution landscape topology is symbolled by a square with the corresponding ID. The widths of the edges are positively correlated with the transformations between the linked landscape topologies, in 100 randomized parameter sets. (C) Transformations induced by *r*_*x*_ . As *r*_*x*_ increases, the transformation from one solution landscape topology to another is indicated by arrows, with the arrow widths positively correlated with the count of transformation in 100 randomized parameter sets. (D) Relationship between the hyper parameters and their directions of cell fate changes. For example, with the increase of hyper parameter *P*_*x*_*up*_, the steady state of the solution landscape bias towards the direction of S2 and LX. So as the other hyper parameters. (E) Categorization of parameters into the four classes of hyper-parameters.

Before introducing the construction of HSL, it is worth to introduce the simplified HSL, an analytical method based on Morse inequalities (Methods). Morse inequality establishes a fundamental relationship between stationary points and manifolds, specifically that the difference between the number of even-order and odd-order saddle points equals one for the manifolds studied here. This relationship persists through local bifurcations, constraining the possible transitions between different landscape topologies. Furthermore, we observed that local bifurcations without limit cycles (such as saddle-node and pitchfork bifurcations) follow a consistent pattern: they typically involve the simultaneous creation or annihilation of adjacent-index saddle points: index−*k* saddle point and index−*k* ± 1 saddle point simultaneously. This feature offers a generation rule of simplified HSL (Methods and Figure S2), instructed by which we can generate it by a process of adding or deleting nodes and edges.

The practical algorithm for constructing the HSL is mainly based on sampling and bifurcation analysis (Figure S1 and Methods). Given that the HSL is inherently a graph composed of nodes and edges, the algorithm is divided into two main components: identifying the nodes and connecting them via edges. The node identification process involves several steps: sampling parameter sets, constructing the solution landscape for each set, and clustering these landscapes. Enhanced sampling may be necessary to ensure node completeness. Notably, prior knowledge, such as the simplified HSL derived from Morse inequality, can aid in node identification during clustering. For the CIS network, we further integrate prior knowledge to exclude biologically-unrealistic landscapes (e.g., those with limit cycles). Edge connection calculates bifurcations by varying one parameter at a time — driving transitions between topologies (e.g., from ID 1 to ID 12). If new nodes emerge, the process iterates with enhanced sampling; otherwise, the HSL is finalized. Since bifurcations are induced by single parameters, edges represent co-dimension 1 types (e.g., saddle-node).

### HSL Enables Systematic Analysis of Parameter-Dependent Bifurcations

The HSL framework provides comprehensive insights into cell fate transition behaviors. Extensive bifurcation analysis across all 16 parameters using randomized sets revealed more frequent bifurcations between common, low-complexity landscapes, reflecting the prevalence of simpler landscape types in parameter space. (Figure 2B). Besides, the solution landscape topology labeled ID 13 emerges as the most highly connected node in the HSL. This topology, representing a bistable landscape with two differentiated states (LX and LY) separated by a saddle point, appears to be the most probable landscape of the CIS motif, as evidenced by its frequent connections to other topologies (Figure S2).

Parameter changes can trigger sequential transformations between solution landscapes, represented as directional routes in the HSL. For example, increasing the parameter *r*_*x*_ produces a systematic progression through different landscape topologies. In this progression, certain topologies act as “sources” (ID41 and ID23, with multiple outgoing edges) or “sinks” (ID2, with multiple incoming edges), revealing how parameter changes can systematically eliminate certain attractors and bias the system toward specific cell fates (Figure 2C).

This hyperparameter framework provides a simplified yet powerful way to understand how different parameters influence cell fate decisions. While a given parameter can induce different bifurcation types depending on the initial landscape, it often produces consistent functional outcomes, such as the annihilation of an LY attractor or the creation of an LX attractor (Figure S10, Section 7 in SI). We therefore classify parameters into distinct hyperparameters based on these shared functional effects. Analysis of parameter-induced changes revealed that the 16 parameters naturally cluster into four categories, characterized by their effects on X and Y gene expression levels (Figure 2D and Figure S2). We formalized these categories as hyperparameters: *P*_*x*_*up*_, *P*_*y*_*up*_, *P*_*x*_*down*_, *P*_*y*_*down*_ (Figure 2D-E). This categorization emerges directly from equation (1) by analyzing how *dx*/*dt* responds to parameter increases. For instance, parameters *r*_*x*_ and *r*_1_, which represent the basal expression and self-activation of gene X respectively, belong to the *P*_*x*_*up*_ category as their increases both elevate X gene expression.

### Different routes in HSL induce distinct cell fate decision behaviors

The HSL provides a global framework for understanding signal-driven cell fate decisions. Each cell fate transition corresponds to a transformation between nodes in HSL, which can occur either through direct parameter changes (“direct route,” blue in Figure 3A) or through a series of sequential bifurcations along HSL edges (“gradual route,” red in Figure 3A). Using the differentiation of uncommitted progenitor cells (transition from landscape ID 21 to ID 13) as a model system, we discovered that different routes between the same initial and final landscapes can produce different cell fate outcomes.

**Figure 3.**
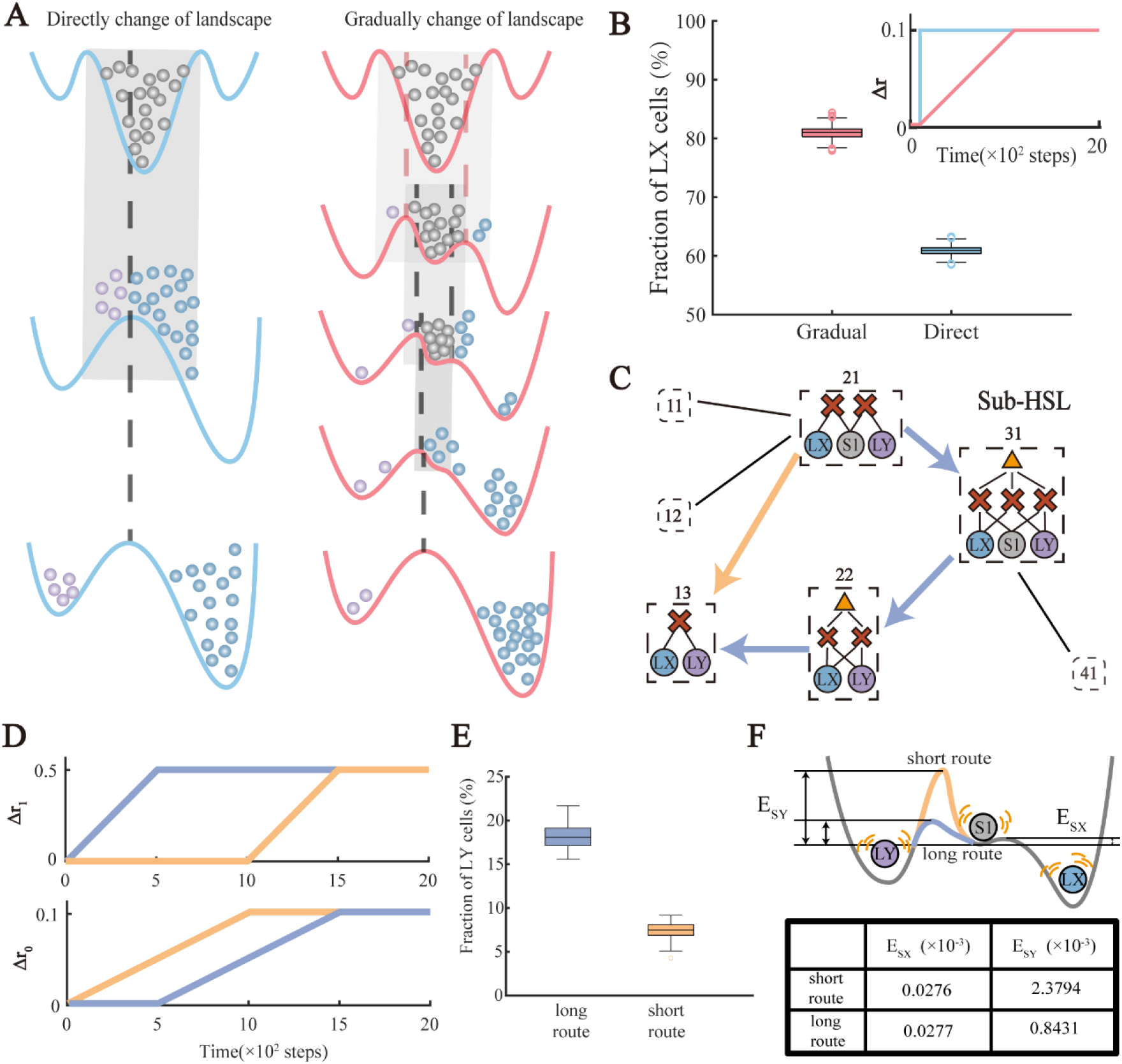
Different routes in HSL induce different cell fate decision behaviors. (A) The schematics of directly change and gradually changes in landscapes. The symbols are the same as the Figure 1. The gray shadows indicate the boundary of the uncommitted progenitor cell attractors in the upper landscape. The black dash lines indicate the position of the saddle points in the bottom landscape. (B) The fraction of LX cells induced by direct (blue)and gradual routes(red), respectively. The parameter changes from the initial to the final in these two routes were shown in the upper box. Simulations were performed with n=1,000 cells for each condition. In the gradual route, parameters changes by the following rules: *r*_1_(*t*) = *r*_1_(0) + 5.5 * Δr(*t*), *r*_4_(*t*) = *r*_4_(0) + 5 * Δr(*t*), *r*_*x*_ (*t*) = *r*_*x*_ (0) + 1.1 * Δr(*t*), *r*_*y*_(*t*) = *r*_*y*_ (0) + Δr(*t*). The noise level in simulations is set to 0.02. Simulations in each route are repeated for 1,000 times to calculate the standard deviation. The standard deviation is 7.6 for direct route and 10.0 for gradual route. (C) Long route and short route in HSL. In the subset of HSL, the short and the long routes from ID 21 to ID 13 solution landscape topologies are indicated by yellow and blue arrows, respectively. (D) Changes in parameters over time for different cell fate decision routes. Blue and yellow lines correspond to the long and the short routes, respectively. The routes correspond to alternatively increase the basal expression level Δ*r*_0_(*t*) and the self-activation level Δ*r*_1_(*t*). The parameters are changed as the following rules: *r*_1_(*t*) = *r*_1_(0) + 1.1 * Δ*r*_1_(*t*), *r*_4_(*t*) = *r*_4_(0) + Δ*r*_1_(*t*), *r*_*x*_ (*t*) = *r*_*x*_ (0) + 1.1 * Δ*r*_0_(*t*), *r*_*y*_ (*t*) = *r*_*y*_(0) + Δ*r*_0_(*t*). (E) The fraction of LY cells induced by different routes. The correspondence between colors and routes is the same as (D). Simulations were performed with n=1,000 cells for each condition. The noise level in simulations is set to 0.02. Simulations in each route were repeated for 100 times to calculate the standard deviation. The standard deviation is 15.9 for long route and 9.2 for short route. (F) The schematic of the landscape near bifurcation. Symbols are the same as the Figure1. The energy barrier of different route is shown in the corresponding color in (E). The energy barrier calculated by GMAM is shown in the table.

To investigate these route-dependent behaviors, we conducted simulations modulating four key parameters: basal expression levels (*r*_*x*_, *r*_*y*_ ) and self-activation levels (*r*_1_, *r*_4_) of genes X and Y through a signal Δ*r*. To introduce a realistic fate bias, we set the parameter changes for the X lineage 1.1 times stronger than those for the Y lineage (details in Methods).

Our first key finding reveals that the gradual route promotes more efficient differentiation toward the LX fate compared to the direct route (Figure 3B). In stochastic simulations starting with 1,000 cells in the S1 attraction basin, the gradual route produced 30% more LX cells than the direct route under identical noise conditions. This difference appears to arise from distinct decision-making mechanisms: in the direct route, a cell’s final fate strongly correlates with its initial gene expression state, analiogous to direct lineage commitment. In contrast, the gradual route allows fate decisions to evolve with the changing landscape, similar to passing through a priming stage before lineage commitment. Consistent with a recent study from Chen *et al*.^54^, slower rates of landscape change in the gradual route increase the final LX fraction (Figure S7).

Furthermore, the HSL framework reveals multiple gradual routes between the same landscapes (Figure 3C): a short route (ID21→ID13) and a long route (ID21→ID31→ID22→ID13). These routes can be implemented by alternating increases in basal expression (Δ*r*_0_) and self-activation levels (Δ*r*_1_) of both genes (Figure 3D): the long route prioritizes increasing Δ*r*_1_ followed by Δ*r*_0_, while the short route reverses this sequence. From the mathematical perspective, the difference of the two routes is the appearance of an unstable attractor and index-1 saddle. Although the bifurcation between index-1 and index-2 saddle does not produce a new cell state (stable attractor), it influences basin connectivity between attractors, potentially leading to observable changes in cell fate switching probabilities. For example, the ID31 are also called fully-connected stage, in which the attractors are connected by each other, and we consider this special stage may correspond to the priming stage of differentiation^43^.

Despite sharing initial and final landscapes, these routes produce substantially different outcomes: simulations with 1,000 uncommitted progenitor cells showed that the long route generated 2.5 times more LY cells than the short route (Figure 3E). This route-dependent bias can be explained through calculating energy barriers (i.e. the quasi-potential between the saddles and the attractors at each time step calculated by GMAM). By comparing the energy barriers from S1 to LY (*E*_*SY*_ ) and S1 to LX (*E*_*SX*_) along both routes, we found that during the critical decision period near the bifurcation point, when the *E*_*SX*_ is sufficient small (lower than 1 × 10^−4^ here), the long route presents an *E*_*SY*_ approximately three times lower than the short route (Figure 3F and Figure S3). This reduced barrier makes the LY fate more accessible to uncommitted progenitor cells following the long route, demonstrating how route choice can influence cell fate outcomes even when starting and ending points remain the same.

We also noted that the different noise conditions influence cell state transitions, as suggested by Coomer at al.^23^ (Table S11 and S12). In the low-noise conditions, the gradual route and direct route model showed minimal sensitivity to multiplicative noise types (*σ* * *x* * *dW*_*t*_, *σ* * (1 + *x*) * *dW*_*t*_, *σ* * (1/*x* ) * *dW*_*t*_), as deterministic mechanisms dominate. In contrast, the long and short routes showed significant sensitivity to noise structure (Table S11). Under high-noise conditions, all routes converged to identical steady-state distributions regardless of noise type, as stochastic forces overwhelmed pathway-specific landscapes (Table S12). Thus, both noise type and magnitude can affect fate transition dynamics, particularly in barrier-dependent systems like the long and short routes.

### Utilizing HSL to Design the Route of Cell Fate Decision

The HSL provides a comprehensive framework for designing signal-driven cell fate transitions. Consider the differentiation from uncommitted progenitor cells to lineage-X (transition from landscape ID 23 to ID 2), where HSL reveals two distinct routes: a “progression route” (red dashed line, Figure 4A) and an “accuracy route” (blue dashed line, Figure 4A), each with unique fate decision characteristics^43^. In the progression route, the uncommitted progenitor cell attractor disappears first, forcing cells to exit the uncommitted progenitor cell state and differentiate (Figure 4B). Such progression routes may occur when a substantial replenishment of cells is required, for instance, in the hematopoiesis. The hematopoietic system experiences a high rate of cellular apoptosis on a daily basis, necessitating rapid differentiation, regardless of accuracy, to replenish these lost cells. In contrast, the accuracy route first eliminates the antagonistic LY fate, restricting cell fate choices to either maintaining stemness or differentiating specifically to LX, thus ensuring higher differentiation precision. Such accuracy routes may occur in embryonic development, in which accurate differentiation into the expected cell type is emphasized.

**Figure 4.**
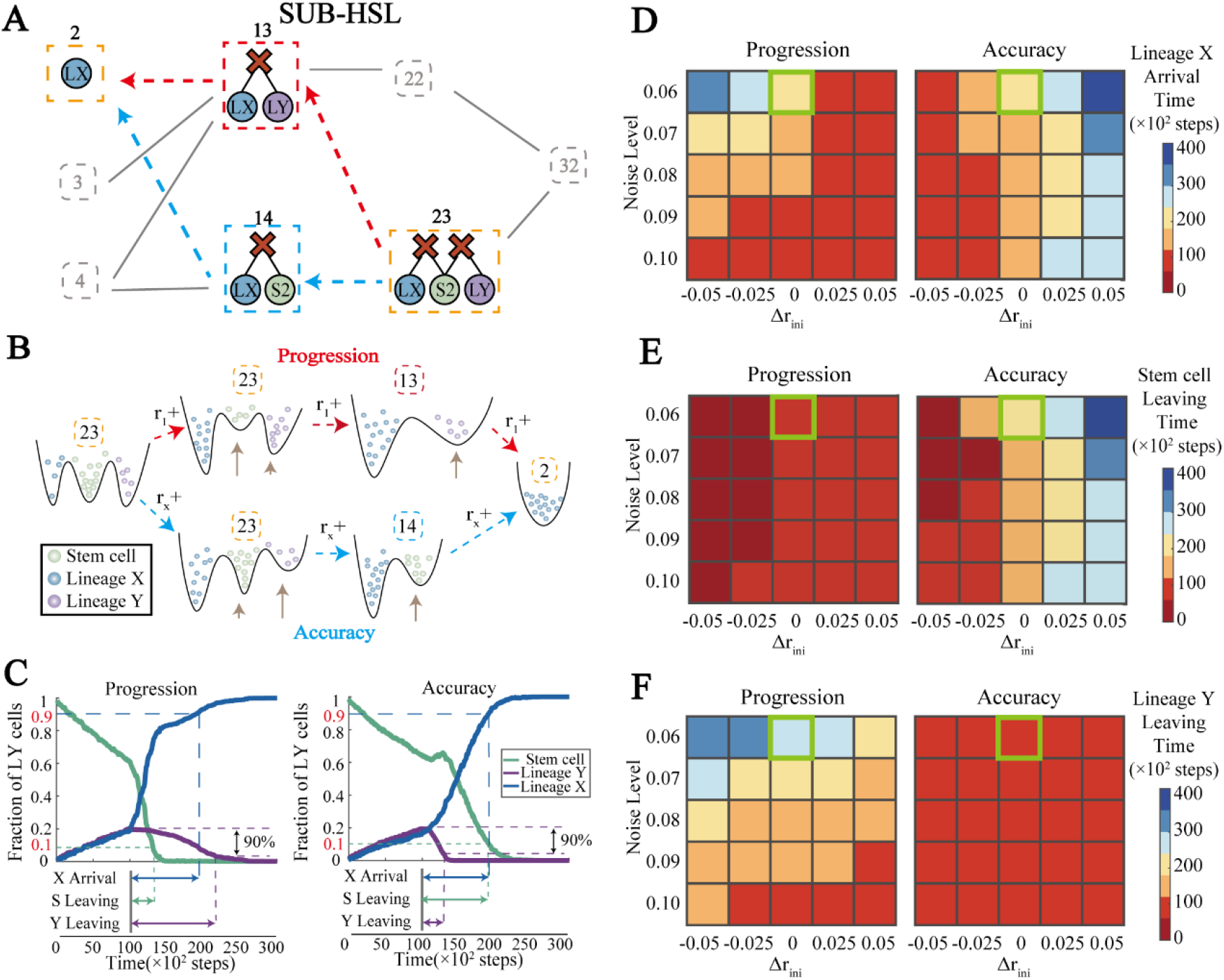
Targeted design of the cell fate transition routes by HSL. (A) Designs of the progression route and accuracy route for differentiation in the HSL of Figure 2(A). We set the initial landscape as ID 23 and the final landscape as ID 2, and there are two routes connecting them. The red arrows mark the “progression route” and the blue arrows mark the “accuracy route”. (B) Schematic changes of the landscapes for the two routes in (A). Along the progression route, with the increasing of *r*_1_, the uncommitted progenitor cell fate disappears first, then the LY fate disappears. Along the accuracy route, with the increasing of *r*_*x*_, the LY fate disappears first, then the uncommitted progenitor cell fate. The green, blue, and purple balls stand for the cells in uncommitted progenitor cell, lineage X, and lineage Y fates, respectively. (C) Changes of the percentage of cell types along the progression (left panel) and accuracy (right panel) routes. Simulations were performed with n=1,000 cells for each condition. The green, blue and purple lines stand for the cells in uncommitted progenitor cell, lineage X and lineage Y states respectively. The noise level and initial parameter is set as indicated by the green box in (D) and (E). (D-F) The heatmap of lineage X arrival time (D), Uncommitted progenitor cell leaving time (E) and lineage Y leaving time (F) in different noise levels and initial parameters for progression and accuracy routes. The initial parameter is set as following: 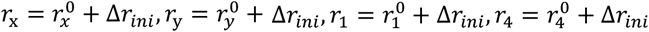. The color scales represent time in simulation steps on a linear scale.

These theoretical routes can be implemented through specific parameter modifications. Based on our analysis in Figure 2, the progression route can be achieved through linear-like increases in basal expression rate *r*_1_, while the accuracy route requires increases in self-activation level *r*_*x*_ (Figure 4B, Figure S4). To compare these routes quantitatively, we conducted stochastic simulations starting with 1,000 cells in the S2 state. After initial equilibration, we tracked the population dynamics along both routes, recording the number of cells in S2, LX, and LY cell type (details in Methods and SI) over time (Figure 4C).

To characterize route efficiency, we defined three key temporal indicators (Figure 4C). The lineage X arrival time measures when the cell type LX population exceeds 90%, providing a metric for successful differentiation. The lineage Y leaving time indicates when the cell type LY population decreases below 90% of its maximum, representing clearance of the alternative fate. Finally, the uncommitted progenitor cell leaving time marks when the uncommitted progenitor cell (S2) population drops below 10%, indicating exit from the uncommitted progenitor cell state. According to their definitions, these three metrics characterize the temporal aspects of different fate-determination processes by statistically simulating the collective behavior of various cell types. These temporal indicators are crucial for systems such as hematopoiesis. For instance, a recent study showed rapid depletion of the common myeloid progenitor (CMP) pool to accelerate downstream differentiation during polymicrobial sepsis-induced inflammation^55^.

These metrics revealed distinct rate-limiting steps for each route. In the progression route, rapid uncommitted progenitor cell exit precedes other transitions, with lineage X arrival time closely matching lineage Y leaving time—indicating that exit from lineage Y is rate-limiting. Conversely, in the accuracy route, lineage X arrival closely tracks uncommitted progenitor cell leaving time, suggesting that uncommitted progenitor cell exit is the primary bottleneck.

Further analysis across varying noise levels and initial landscapes revealed environment-dependent route efficiencies. Initial landscape geometry, controlled by parameter shift Δ*r*_*ini*_, particularly affects energy barriers between uncommitted progenitor and differentiated states (Figure S4). Heat map analysis of temporal indicators across these conditions (Figure 4D-F) revealed several key insights into route behavior. The overall efficiency of each route, measured by lineage X arrival time, strongly correlates with its characteristic rate-limiting step. Under conditions of low noise and low energy barriers, the progression route shows diminished performance as cells have difficulty exiting the LY state. In contrast, the accuracy route faces its greatest challenges at low noise levels combined with high energy barriers, where cells exhibit delayed exit from the uncommitted progenitor cell state, resulting in extended differentiation times. These findings demonstrate how HSL can guide the design of effective differentiation strategies by identifying and addressing route-specific bottlenecks in cell fate transitions.

### The Application of HSL in Seesaw Model

The HSL framework can be extended to various network structures. A particularly significant application lies in the “seesaw” model of cellular reprogramming (Figure 5A), which demonstrates how maintaining regulatory balance is crucial for restoring pluripotency^56^. In this model (details in Methods), pluripotency factors OCT4 and SOX2 activate each other while promoting their respective lineages, while lineage-specifying gene groups MEs and ECTs exhibit mutual antagonism and suppress pluripotency (Figure 5A). We constructed the HSL using eight key parameters from the original model: external signals that either induce ( *C*_0_, *C*_*S*_, *C*_*M*_, *C*_*E*_ ) or inhibit ( *I*_*O*_, *I*_*S*_, *I*_*M*_, *I*_*E*_ ) the expression of OCT4, SOX2, MEs, and ECTs, respectively.

**Figure 5.**
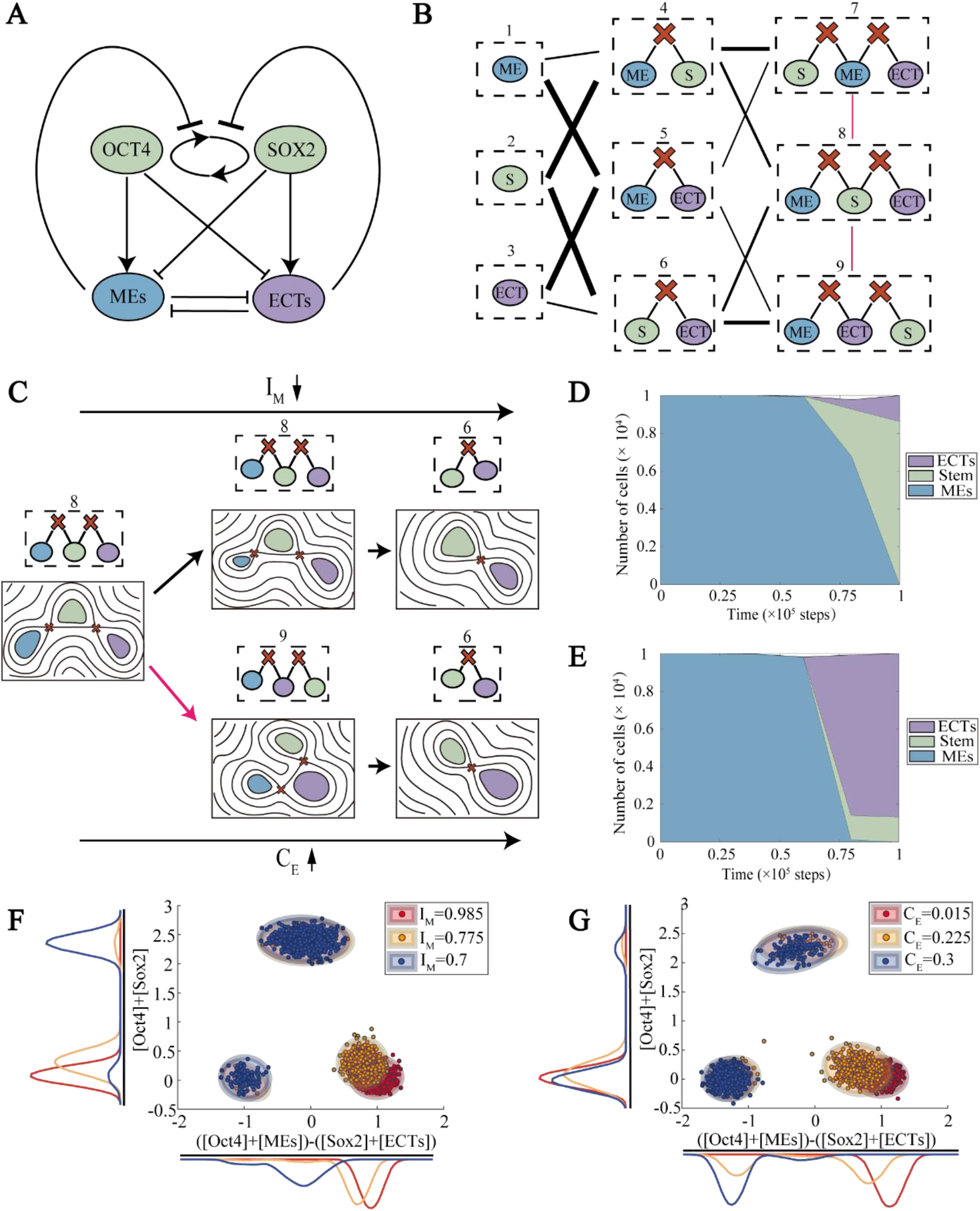
HSL of the seesaw model reveals how different routes influences fate distributions. (A) The topology of the gene regulatory network underling the original seesaw model^56^. Color schemes and the symbols are the same as Figure1(A). (B) The HSL with weighted edges for seesaw model. Each solution landscape topology is symbolled by a square with the corresponding ID. The symbols are the same as Figure2(A). The widths of the edges are positively correlated with the transformations between the linked landscape topologies, in 100 randomized parameter sets. The pink lines between ID 8 and ID 9 (ID 7) represent heteroclinic flip bifurcations. The cell types are distinguished by the stemness and the bias (Methods). (C) Schematics on how different parameter changes induce different landscape transformations, forming different routes. The upper route corresponds to the change of solution landscape topologies from ID 8 to ID 6, induced by the *I*_*M*_ linear decreasing over time. The bottom route corresponds to the change from ID 8 to ID 9 and then to ID 6, induced by the *C*_*E*_ linear increasing over time. (D-E) Stacked area plots show the distribution of cell types at different times for 8-6 route(D) and 8-9-6 route(E). The green, blue and purple areas stand for the number of cells in stem cell, lineage X, and lineage Y states, respectively. 1,000 cells are set in ME state in ID 8 solution landscape initially and reach the ID 6 solution landscape finally, with different parameters changing linear over time. (F-G) The distribution of cell types at different parameter situation for 8-6 route(F) and 8-9-6 route(G). Dots with different colors represent cells at different parameter situation. The cells are clustered by Gaussian mixture method.

The resulting HSL reveals that two dominant routes in seesaw model: (1) the differentiation: from landscapes ID 5→ID 1, ID 5→ID 3 and (2) reprogramming: from landscapes ID 4→ID 2, ID 6→ID 2 (Figure 5B, Figure S5). Starting from the ID 8 landscape, as the same initial landscape in the reference^56^, transitions predominantly target ID 4/6 landscapes (reprogramming routes), aligning with the model’s major focus. However, even between the same initial (ID 8) and intermediate (ID 6) landscapes, different parameter modifications can induce distinct reprogramming behaviors.

When ME fate suppression dominates (through increased *I*_*M*_ ), the landscape transforms directly from ID 8 to ID 6. However, when ECT fate activation prevails (through increased *C*_*E*_), the transition proceeds via an intermediate ID 9 landscape (Figure 5C). Unlike prior bifurcations, the ID 8-to-ID 9 shift is a heteroclinic flip bifurcation, altering connectivity without changing fixed-point numbers (Figures 5C and S5). And when heteroclinic bifurcations redirect trajectories to ID 9, subsequent transitions overwhelmingly favor ID 6 (Figure 5B). These distinct routes produce markedly different fate outcomes: ME suppression forces direct reprogramming to stem cells (Figure 5D and 5F), while ECT activation enables both reprogramming to stem cells and transdifferentiation to ECT cells (Figure 5E and 5G).

The fate heterogeneity arise from different topological connections between attractors. In the ID 9 landscape, ME cells connect to ECT cells through an index-1 saddle, while in ID 8, ME cells connect directly to the stem cell state. As signaling increases, the ME attractor gradually destabilizes, enhancing transition probability to adjacent states according to the landscape’s topology.

This application of HSL to the seesaw model not only demonstrates the framework’s versatility but also reveals how different routes through the landscape can fundamentally alter cell fate outcomes. These insights suggest that careful consideration of transition routes, not just initial and final states, is crucial for optimal control of cell fate engineering.

## Discussion

The quest to understand cell fate decisions through landscape perspectives has evolved greatly since Waddington’s initial metaphor. We construct a holistic map of cell fate decision by applying a HSL approach. HSL incorporates fundamental principles of landscape transitions, particularly how local bifurcations involve the simultaneous creation and annihilation of adjacent-order stationary points. Through systematic analysis of the CIS network, we identified hub nodes within the HSL and consolidated the numerous parameters into four key hyperparameters that characterize directional landscape changes. This framework demonstrates how different routes through the landscape can shape cell fate heterogeneity, providing a global perspective for designing cell fate transitions with specific properties, such as optimizing for either progression efficiency or fate accuracy.

As the node of HSL, the solution landscape approach offers several advantages over traditional methods. Unlike energy landscapes constructed by solving Fokker-Planck equations^14^ or transition path optimization^30^, it efficiently resolves the complete topological hierarchy of stationary points (attractors, 1-saddles, 2-saddles, etc.) and their connections through saddle dynamics. This topology-first strategy eliminates the need for prior state definitions and extensive stochastic simulations — particularly critical in high-dimensional gene regulatory networks. Such strategy of connecting stationary points has gained much attention in biological network modeling^36^, such as Directed Acyclic Graph (DAG) that shares conceptual similarities with solution landscape, and Bifurcation Diagrams (BD) share conceptual connections with HSL. Building on these, the solution landscape serves as a computable tool for searching points and connections via saddle dynamics, while HSL extends it to ensembles of landscapes and bifurcations in gradient and non-gradient systems. Key differences from BDs in geometric approaches include: (1) basic structures (solution landscapes vs. “MS components”); (2) HSL’s focus on node connectivity (transition pathways) versus BDs’ emphasis on topological boundaries; and (3) HSL’s multi-layered hierarchy versus BDs’ continuum representation. For instance, Figure 1E depicts a “dual cusp” in geometric approaches, and restricting parameters *r*_*x*_ and *r*_*y*_ to the range of 0.1-0.25 yields a consistent sub-HSL (Table S9 and Figure S6).

Building on these strengths, HSL provides a unified framework for investigating both noise-driven and signal-driven cell fate decisions at distinct analytical scales. At the node level, each solution landscape represents a static landscape, enabling the study of noise-driven effects through the extraction of transition states and transfer pathways. At the edge level, connections between solution landscapes reflect landscape changes associated with bifurcations, facilitating exploration of signal-driven behaviors and cellular adaptation to environmental stimuli. Overall, the HSL enhances our understanding of the intricate interplay between noise and signals in cellular fate decision, positioning it as a valuable tool for advancing insights into complex biological processes.

The practical utility of HSL lies in its ability to guide the design of cell fate decision routes. Acting as a holistic map, it enables identification of optimal paths to desired cell states. With advancing capabilities in synthetic biology^57–59^, these theoretically designed routes can be experimentally implemented through timed induction of transcriptional regulators. Our analysis of progression and accuracy routes revealed distinct rate-limiting steps, providing crucial insights for route design in real systems. Future applications could extend to optimizing pathways under multiple constraints to meet diverse functional requirements.

In summary, the HSL framework advances our understanding of cell fate decisions by providing a holistic, topology-driven map that unifies noise- and signal-driven transitions while overcoming parameter dependencies. It equips researchers with powerful tools for rationally designing cellular engineering strategies, from optimizing differentiation routes to precise reprogramming. As synthetic biology advances, HSL’s applications promise to extend beyond cells to diverse dynamical systems, catalyzing breakthroughs in biology and beyond.

### Limitations of the study

While this work establishes HSL as a powerful framework, several challenges remain for future investigation.

Firstly, the theory and algorithms of HSL need further refinement for application to more complex systems, particularly those exhibiting dynamic behaviors like limit cycles^32^. The HSL framework currently relies on saddle dynamics to trace stationary points and their connections, which inherently cannot detect limit cycles as the algorithm cannot converge to limit cycles. While this restricts our current analysis of periodic solutions, we fully acknowledge the critical need for advanced algorithms to efficiently identify limit cycles and characterize their relationships with stationary points. Future work will focus on extending the HSL methodology to incorporate cycle-oriented dynamics, enabling systematic exploration of solution landscapes involving both steady states and oscillatory behaviors.

Secondly, extending the HSL framework to high-dimensional gene regulatory networks^46,60^ introduces substantial computational challenges, due to the exponential increase of stationary points and the complexity of bifurcations in the HSL constructions. To mitigate this, we prioritize biologically informed dimensionality reduction—for example, focusing on core regulatory parameters (Oct4/Sox2/Nanog in pluripotency networks) and aggregating redundant nodes. Future integration with parallel computing will further expand the scope to complex high-dimensional systems.

Thirdly, the high co-dimension bifurcations need to be explored. In the process of constructing HSL, the change of solution landscape was induced by one single parameter. It means that the co-dimension of bifurcations in HSL equals 1 and the bifurcations are mostly saddle-node or flip. Though we demonstrate a co-dimension 2 bifurcation (dual cusp) in SI, the high co-dimension bifurcations induced by multiple parameters are still needs exploration. And how to identify the bifurcation type directly from HSL is also a direction worth exploring. The HSL is constructed through single-parameter variation, meaning that only co-dimension 1 bifurcations (e.g., saddle-node, flip) are captured. As such, bifurcation types are not directly labeled in the HSL. Future work using multi-parameter analysis will allow identification of higher-order bifurcations and enable more refined bifurcation classification within the HSL.

## RESOURCE AVAILABILITY

### Lead Contact

Further information and requests for resources should be directed to and will be fulfilled by the Lead Contact, Lei Zhang.

### Materials Availability

This study did not generate new materials.

### Data and Code Availability

All data reported in this paper will be shared by the lead contact upon request.

All original code has been deposited at Zenodo and is publicly available as of the date of publication. DOIs are listed in the key resources table.

Any additional information required to reanalyze the data reported in this paper is available from the lead contact upon request.

## ACKNOWLEDGMENTS

We would like to thank J. Yin and H. Su for their advice on mathematical methods, and N. Yang. and Y. Yu. for their support in theoretical concepts. We also extend our gratitude to all members of Lei’s group for their support and assistance. L.Z. was supported by the National Key Research and Development Program of China (No. 2024YFA0919500) and the National Natural Science Foundation of China (No. 12225102, T2321001, 12288101, 12426653). Z.L. was supported by the National Key Research and Development Program of China (No. 2021YFA0910700, 2024YFA0919500), and Fundamental and Interdisciplinary Disciplines Breakthrough Plan of the Ministry of Education of China (JYB2025XDXM502). This work supported in part by Peking-Tsinghua Center for Life Sciences.

## AUTHOR CONTRIBUTIONS

X.Z., Z.L. and L.Z. conceived the project; X.Z. performed the analysis and computational simulations; Z.L. and L.Z. supervised the project; X.Z., Z.L. and L.Z. wrote the paper.

## STAR★METHODS

### Key Resources Table

| REAGENT OR RESOURCE | SOURCE | IDENTIFIER |
| --- | --- | --- |
| Software and algorithms |  |  |
| MATLAB r2022b | The Mathworks Inc. | https://matlab.mathworks.com/ |
| GHiSD (Generalized High-index Saddle Dynamics) | (Yin et al., 2021)<sup>42</sup> | N/A |
| hsl-project | This paper | Zenodo: 10.5281/zenodo.18006147<br>GitHub:<br>https://github.com/zzzhangxy98/hsl-project.git |

## METHOD DETAILS

### Gene Regulatory Network

The GRN used in this work follows previous models^43,61^. The equation (1) derives from a biochemical reaction model incorporating dimerization, reversible DNA binding, transcription, translation, decay, and ectopic overexpression^61^. The original 14-dimensional system was reduced to equation (1) using quasi-steady state assumptions.

### Generalized High-index Saddle Dynamics (GHiSD)

k-GHiSD identifies index-k saddle points in dynamical systems^42,62^, represented by dynamics (3). This method can be interpreted as a reverting evolution along the unstable subspace *𝒲*^u^(***x***):

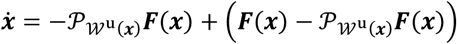

Where 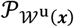 stands for the orthogonal projection operator onto the unstable subspace *𝒲*^u^(***x***). The second equation for the saddle dynamic aims to find an orthonormal basis {***v***_**1**_, …, ***v***_***k***_} of the unstable subspace ***W***^***u***^(***x***).

### Algorithm for constructions of the solution landscape

Generally speaking, we can use the downward-search and the upward-search algorithms to construct the solution landscape^41,44^, which starting from the *k*-index saddle points and the stable points respectively.

Downward search

1. Given an *k*-index saddle ***x***, calculate the unstable directions{***v***_1_, …, ***v***_*k*_}.
2. Choose an unstable direction ***v***_***j***_ and perturb the initial state along this direction.
3. Start from the point ***x*** ± ***εv***_***j***_, find the *m*-index saddle point (*m* < *k*) using the m-GHiSD.
4. Record the new saddle point, and the connection relationship.
5. Repeat the process for all of the unstable directions and all of the indices until no new saddles can be found.

Upward search:

1. Given an *k*-index saddle ***x***, calculate the stable directions{***v***_1_, …, ***v***_*p*_}.(suppose the highest index of saddle points is *k* + *p*)
2. Choose a stable direction ***v***_***j***_ and perturb the initial state along this direction.
3. Start from the point ***x*** ± ***εv***_***j***_, find the *m*-index saddle point (*m* > *k*) using the m-GHiSD.
4. Record the new saddle point, and the connection relationship.
5. Repeat the process for all of the stable directions and all of the indices until find the highest index saddles.

In practice, we use the upward search and the downward search algorithm together.

### Morse inequality

Morse theory is a geometrical theory that studies the topological relationship between stationary points and manifolds. For the dynamical system we studied, it satisfies the constraints in reference^63^ and the Morse inequalities for the dynamical system can be shown by the following expression:

Let *α*_*i*_ be the number of stationary points of index *i* and *β*_*i*_ be the *i*-th Betti number (i.e., the rank of the *i*-th homology group). We have the following inequalities:

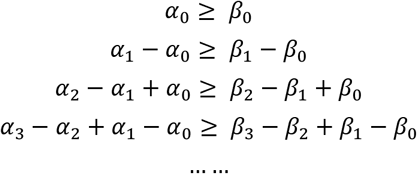

The Betti numbers for ℝ^2^ are as follows: *β*_0_ = 1, *β*_*i*_ = 0 for *i* ≥ 1.

In the context of a two-dimensional system, the maximum index of saddles is 2, indicating that there are no saddles of index 3 or higher:*α*_***j***_ = 0 *for* ***j*** ≥ 3.

Therefore, the inequalities can be simplified to the following:

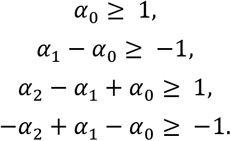

The last inequality can be rewritten as:

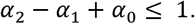

Considering the last two inequalities together, we find that the number of saddles satisfies:

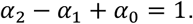

Since bifurcations do not alter the topological features of the manifold, this equation holds both before and after bifurcations. In our study, we observe that bifurcations affect the number of stationary points according to the following rules: *k*-index and (*k* ± 1)-index saddles appear and disappear simultaneously.

### A framework of constructing simplified HSL

We present a theoretical framework for constructing simplified hyper solution landscape (simplified HSL), which primarily focuses on the topological structures of the solution landscape (Figure S2). The methodology encompasses the following steps: 1) fixing the starting and ending solution landscapes; 2) applying generation rules derived from Morse inequalities to create the solution landscapes between them; and 3) pruning based on prior knowledge.

In the specific application to the CIS network, our algorithmic process is detailed as follows:

1. Fixing Starting and Ending Points: The simplest solution landscape is characterized by the presence of a single attractor. According to our prior knowledge^64^, the most complex solution landscape for the CIS network consists of a 2-saddle, four 1-saddles, and four attractors (Figure 1C).
2. Generation Rules: Within the CIS network, the methods for expansion include two types: adding a 2-saddle and a 1-saddle to the solution landscape, or adding a 1-saddle and a 0-saddle (attractor) to the solution landscape.
3. Modifications: By integrating prior knowledge of the CIS network, we eliminate structures that obviously correspond to other topologies.

The resulting simplified hyper solution landscape (simplified HSL) can then be utilized in the subsequent construction of the HSL.

### HSL Construction Algorithm

The flowchart for constructing the HSL (Figure S1) is organized into the following steps:

1. Input Data: Begin by inputting the GRNs (or ODEs), along with the corresponding parameter space. The range of parameter was set to [0,10] in our work.
2. Initial Parameter Sampling: Conduct initial sampling of the parameter set using Latin Hypercube Sampling or leveraging prior knowledge, such as a simplified HSL constructed based on Morse inequality.
3. Solution Landscape Generation: For the parameter sets identified in the previous step, generate solution landscapes through both upward and downward searches.
4. Clustering: Cluster the pool of solution landscapes into distinct topologies by the number and type of stationary points.
5. Enhanced Sampling: Perform enhanced sampling for topologies that exhibit insufficient sampling to ensure comprehensive representation.
6. Bifurcation Search: Execute a bifurcation search by generating solution landscapes across a range of parameters to identify critical transitions.
7. Output: If a new topology of solution landscape is discovered, return to step 5 for further enhanced sampling; otherwise, proceed to output the final HSL.

The computational costs for HSL are shown in Table S13 and Table S14.

### The descriptive name for CIS motif

The descriptive names are shown by the following format:

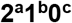

Where **a (a=0**,**1)** is the number of index-2 saddle, **b (b=0**,**1**,**2**,**3**,**4)** is the number of index-1 saddle, and **c (c=S1**,**X**,**Y**,**S2)** is the type of attractors. The descriptive names of different IDs are shown in Table S10.

### Geometric Minimum Action Method (GMAM) and quasi-potential

The GMAM^52,53^ is a numerical algorithm used to compute the Minimum Action Path (MAP) connecting two metastable states in a dynamical system. The key idea behind the GMAM is to reformulate the Freidlin-Wentzell action functional on the space of curves. We apply the GMAM method to calculate the transition pathways between two attractors ***x***_**1**_ and ***x***_**2**_ and the quasi-potential ***V***(***x***_**1**_, ***x***_**2**_) representing the energy barrier.

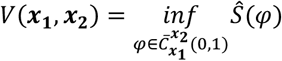

Where 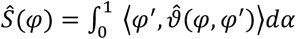 is the action calculated by GMAM.

### Stochastic simulation

We introduce noise into Equation (1) to obtain the Langevin equation^14,30^, expressed as follows:

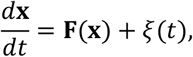

where ***F***(***x***) represents the right-hand side of Equation (1), and *ξ*(*t*) denotes the Gaussian white noise. The properties of the noise are given by:

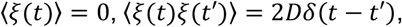

where *D* is the noise level.

### Ranges of bifurcations

The range for CIS motif: the following multiplier to the initial parameters:

{0.0001,0.001,0.01,0.1,0.5,0.9,0.95,0.99,0.995,0.999,1,1.001,1.005,1.01,1.05,1.1,1.5,2,2.5,5,10}

The ranges for seesaw motif: 0-1 with the interval of 0.05.

### The framework of hyperparameter

Our hyperparameter classification methodology is grounded in systematic analysis of solution landscape transitions, as illustrated in Figure 2C and Figure S2C in manuscript. The parameter ranges for bifurcations are shown in the “Ranges of bifurcations” section in Methods. We classify parameters based on the dominant trend of attractor changes induced by their increase:

1. Primary labeling: A parameter is assigned a directional label by the dominate effects (e.g., *P*_*x*_*up*_, *P*_*y*_*down*_), shown by the arrows with orange color in figure S2c. For example, if the dominate effect for a parameter is the disappearance of LY attractors (e.g. the widest arrow in Figure 2C), it is classified as *P*_*x*_*up*_ *or P*_*y*_*down*_ .
2. Final labeling: If a parameter shows mixed effects (with multiple labels, such as the co-occurrence of *P*_*x*_*up*_and *P*_*y*_*down*_), we resolve ambiguity using the parameter’s role in the governing equations (Eq. 1): For example, when a parameter were marked by both *P*_*x*_*up*_ and *P*_*y*_*down*_, if it modulates the *F*_1_(*x, y*) term governing X dynamics (e.g. *r*_*x*_ ), it is designated *P*_*x*_*up*_; otherwise, if it affects *F*_2_(*x, y*) (Y dynamics), it is classified as *P*_*y*_*down*_.

### The equations of seesaw model

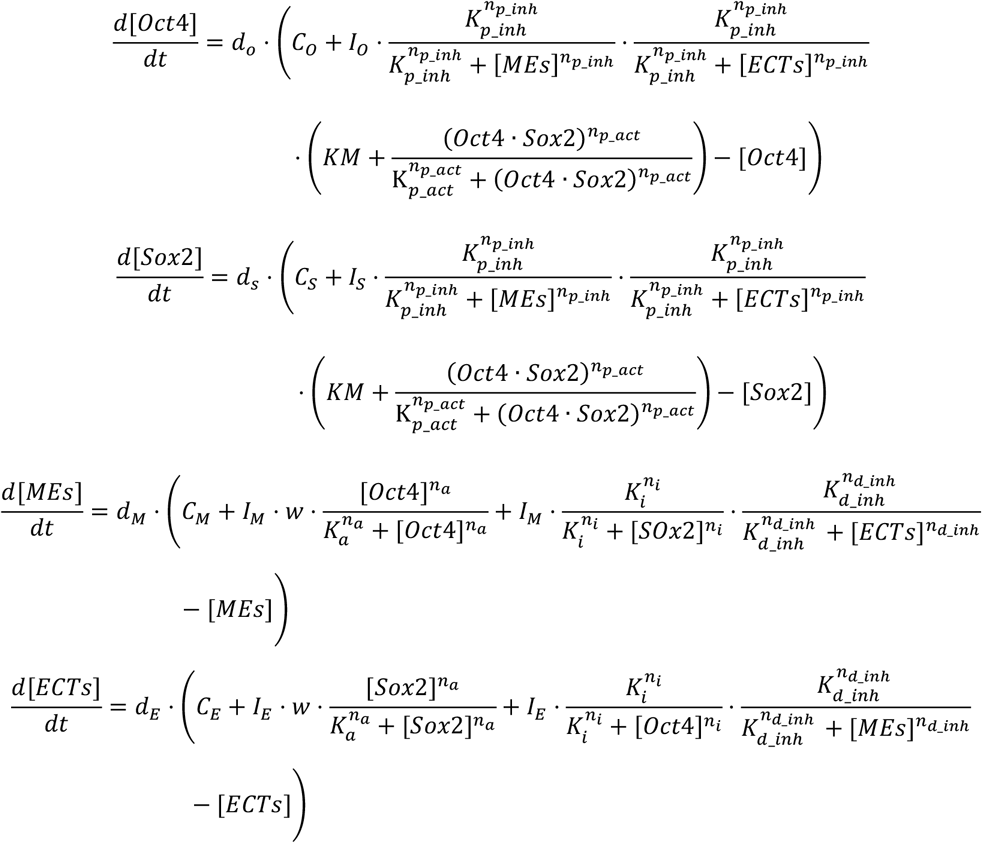

### Classification criteria of cell types

#### For the CIS motif

The attractors are distinguished by the expression level of X and Y gene (Figure 1B). In the simulations, we can use the following three methods to define the cell types:

1. By the fate bias ([*X*] − [*Y*]) of gene expression level. The cell type of uncommitted progenitor cell, LX and LY is corresponding to −0.5 < [*X*] − [*Y*] < 0.5, [*X*] − [*Y*] ≥ 0.5 *and* [*X*] − [*Y*] ≤ −0.5, where the [*X*] and [*Y*] represent for the expression level of gene X and gene Y, respectively.
2. By the hyper spherical basin approximation. Cells are classified by calculating the minimum Euclidean distance to the attractors at each timestep, with a cutoff of 0.5 (dimensionless units) to define basin boundaries. If the distance of a cell and a certain attractor is less than 0.5, we define the cell type as the corresponding attractor.
3. By relaxation-based fate assignment. For each cell at every timestep, we iteratively relax its position to convergence and assign fates based on the terminal attractor.

In our simulation, these methods have the similar results (Figure S8 and S9), so we choose the method 1 to make the Figure4C.

### For the seesaw model

The cell types are distinguished by the stemness (*stemness* = [*Oct*4] + [*Sox*2]) and the bias (*bias* = ([*Oct*4] + [*MEs*]) − ([*Sox*2] + [*ECTs*])). The stem cell state has *stemness* ≥ 1. The ME state and the ECT state corresponds to the case of *stemnes* < 1, *bias* > 0 and *stemnes* < 1, *bias* > 0, respectively.

## SUPPLEMENTAL INFORMATION

**Document S1. Figures S1-S10. Tables S1-S15**.

